# A descriptive study of wetland vegetation, ecology, habitat and Rift Valley fever virus vectors in South Africa

**DOI:** 10.1101/2025.08.13.670234

**Authors:** R. F. Brand, M. K. Rostal, A. Kemp, H Zwiegers, L. de Jager, Z. Gqalaqha, L. van Staden, C.W. van Huyssteen, A. Anyamba, N. Gregory, C. Cordel, W. B. Karesh, J.T. Paweska

## Abstract

Outbreaks of Rift Valley fever (RVF) have been closely linked to environmental conditions that support large vector populations with what is required for breeding of vectors. Fine-scale vegetation surveys were used to assess how wetland plant communities vary across sites with historical RVF outbreaks in Free State Province of South Africa and to detail plant species density that correlates with mosquito habitat.

**Methods:** At 22 sites, we compiled 201 relevés using the modified Braun-Blanquet method in combination with 200 transects using the line-point method developed specifically for this study. A total of 195 plants were identified and vouchered. These data were combined with soil analyses carried out at all 22 sites and with data from bimonthly mosquito trapping at all sites.

**Results and significance:** We identified an increase of invasive plants and upland species compared to the earlier study and observed that drought causes a change in wetland vegetation community structure and species composition. However, the wetland vegetation was still dominated by grasses and sedges, with a limited presence of *Juncus* and forbs. Livestock grazing pressure on wetlands to the drought results in reduced vegetation cover, abundance and diversity. Our work describes a changing landscape that may impact RVFV maintenance and transmission, and provides insights into extensive livestock farming and rainfall over shallow endorheic depressions in temperate grasslands and subtropical riparian environments.

## Introduction

Rift Valley fever virus (RVFV) poses a significant risk to human and livestock health [1–3]. Rift Valley fever (RVF) was first diagnosed in 1930 in western Kenya (Daubney *et al.,* 1931) and has since been identified in West Africa [4–6], East Africa [2, 7–10] and southern Africa, the Middle East and North Africa (MENA) region [3, 11–13] and islands of the Indian Ocean [14]. In southern Africa, RVFV epidemics are typically associated with above-normal summer rains, which produce flooding of suitable wetlands. Such rainfall events frequently occur during the La Niña period of the El Niño–Southern Oscillation (ENSO) cycles [15]. The effect of the ENSO cycles on landscape greening as detected by satellite-derived normalised difference vegetation index (NDVI) has been used to forecast suitable conditions for RVFV epidemics in the region [16–18].

Since the 2010/11 RVF epidemic in South Africa, extensive research has been carried out with various publications investigating links between the 2010-2011 outbreaks and local wetland vegetation and ecology [15, 19, 20]. Brand *et al*., 2018 investigated the relationship between RVF outbreaks in livestock and the phytosociology of wetlands that provide breeding habitat for vector mosquitoes in the genus *Aedes* (Diptera: Culicidae). The floodwater Aedes species, (potential RVF virus vectors) lay eggs on the moist banks off partially-flooded pans and wetlands [17, 21] (Becker 1989). The embryonated eggs survive the winter dry season and, possibly, longer periods of drought (Gjullin & Yates 1946, Horsfall *et al*., 1973). Local rainfall is responsible for flooding of seasonal wetlands, allowing mosquito eggs to hatch. The main RVFV vectors consist of mosquito species in the genus, *Aedes*, particularly in the subgenera, *Aedimorphus* and *Neomelaniconion* [21–23]. Members of the former subgenus appear epidemiologically more significant in the arid areas of West Africa [4, 24], while members of the latter subgenus are more usually implicated in the tropical, subtropical and temperate regions of East and southern Africa [22, 25]. Although not pertinent to the present analysis, the wetlands also attract mosquito species that breed in permanent water bodies such as species of *Culex*, *Mansonia* and *Coquillettidia* (Gregory *et al.*, In prep), some of which serve as horizontal RVFV vectors.

We have continued our research on wetland ecosystems because metrics for vegetation used in RVFV ecology analyses are typically derived at coarse scales (e.g. NDVI) and may not accurately reflect the local conditions to which mosquito vectors complete their life cycle. By conducting fine-scale vegetation surveys at sites previously experiencing RVF outbreaks in South Africa, this work aims to provide a better understanding of, and insights into, the co-occurrence of multi-scale environmental drivers (meteorology, vegetation ecology, wetland categorization, geology, soils) with mosquito relative abundances and RVFV epidemiological findings.

This syntaxonomical/phytosociological study and the previous one of Brand *et al*., (2018), uses theoretical method to analyse the structure and composition of plant communities [26, 27] (Westhoff &, Van Der Maarel, 1978), name syntax using multivariate statistical analysis [28](Hill 1979 identify, and categorize the vegetation composition by ordering into plant families and document changes within and between wetlands and depressions. Various RVF studies detail environmental conditions including meteorology, climatology [15], wetlands, depressions [21] and plants [17] Rossouw *et al.,* 2021), but do not differentiate between wetland categories, or analyse, and describe vegetation to identify plant community structure and composition and compiled syntaxonomical/phytosociological tables.

## Materials and Methods

Our second vegetation ecology study started in October 2016 and was completed in 2018. Eight new sites were added that had not previously been surveyed. These new sites were selected primarily for a cohort sheep study, in which we could monitor the incidence of RVFV seroconversion (Rostal et al., In review). Wetland vegetation surveys were conducted on all 22 sites, using the Braun-Blanquet (B-B) method combined with line-point surveys (L-P). Simultaneously mosquito collections were ongoing, and soil samples were collected for laboratory analysis [19].

The majority of sites surveyed were on working farms with the exception of the SANParks Holpan/Graspan National Park. All wetlands, except the SANParks site were subject to heavy overgrazing, trampling, erosion and soil change. No communal grazing sites were selected for sampling due to the potential damage to instruments, theft, and safety of field staff. An additional consideration was the need for consistency of access to the same plots over several years.

### Study area and site selection

The study area covers approximately 200 × 200 km^2^ (28S−30.45S, 24E−26.65E), the majority of which is in the Free State Province, and some limited areas of Northern Cape province. The study area map (Figure 1; [29], includes wetlands, vegetation units in the Grassland Biome, mostly show in green, which comprises most of the Free State, and in the west, Savanna Biome with its vegetation units, shown in shades of brown.

**Figure 1.**
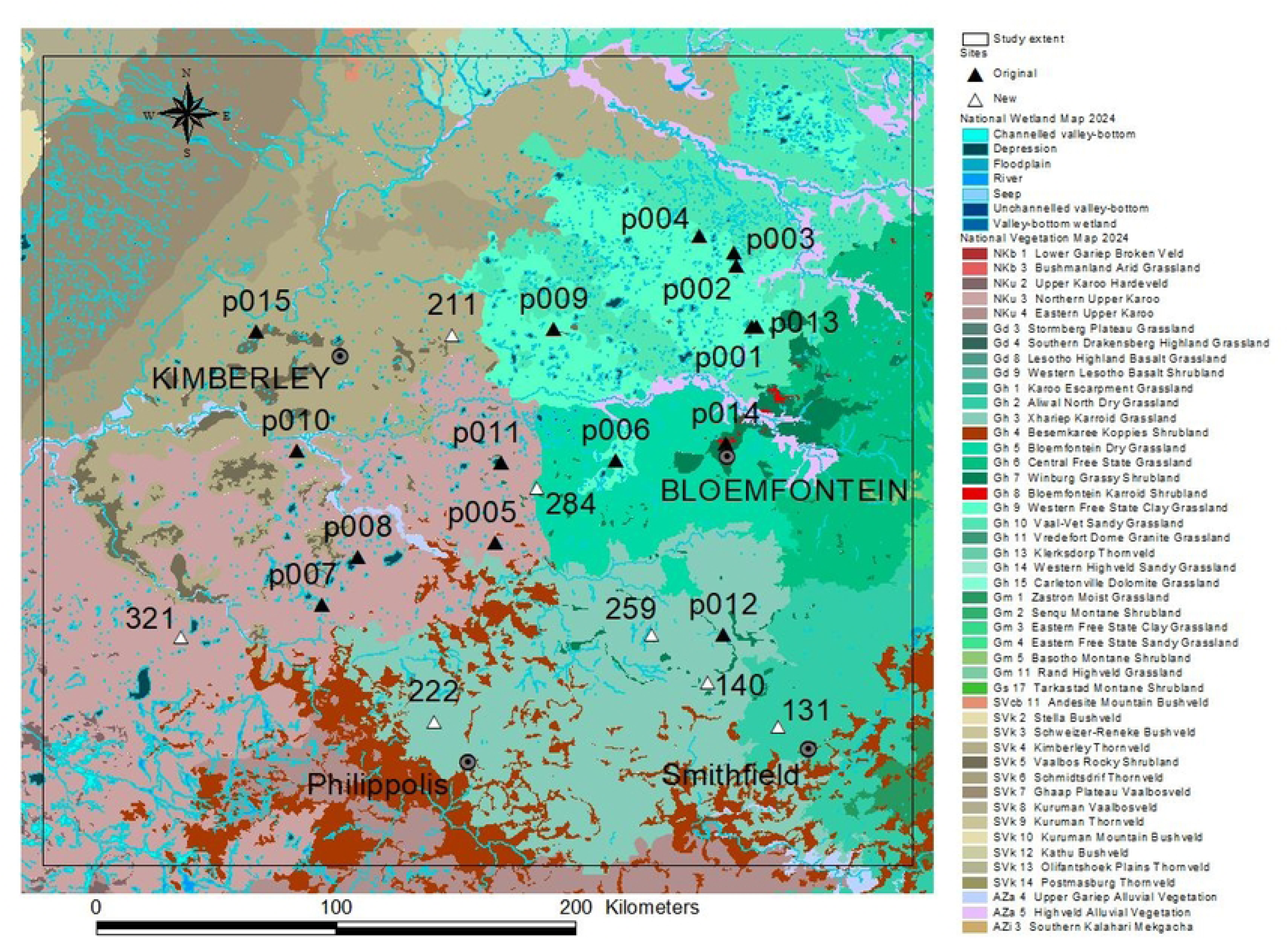
Study sites (triangles) in the Free State and Northern Cape, South Africa, with the initial 15 sites indicated in black, and the seven additional study sites added in 2016 (white). Sites p010 and p015 are in Savanna Biome, all other sites are Grassland Biome. Map courtesy Mr. Leslie Powrie June 2025.

A total of 22 sites were surveyed using both Braun-Blanquet (B-B) and line-point (L-P) methods. This included the original 15 sites. Site selection was primarily focused on assessing the overall ecological conditions of suitable habitats for floodwater mosquito oviposition and larval development, i.e., pans (endorheic wetlands), selected to coincide with livestock farms that had experienced RVF outbreaks, except for the wildlife conservation areas where no RVF outbreaks was recorded [20].

### Geology and Soils

The geology of the area comprises sedimentary rocks of the Karoo Supergroup, with the oldest strata being Dwyka tillites, outcropping in the northwest, overlain by Ecca and younger Beaufort shales and sandstone (Goudie 1991). Dolerite dykes and sills of Stormberg age are numerous, and form the predominant, flat-topped hills characteristic of the landscape. Volcanic andesite outcrops in the far west of the study site comprise the major rock-type at the Holpan/Graspan National Park. The full geology is described in detail by [20].

Detailed soil sampling of 15 sites was carried out in May and June of 2015, using a hand-held soil auger, in the same wetlands where the vegetation and mosquito sampling was done. Subsequent soil collection for the last seven sites was completed in 2017 [30]; supplementary material. The analysis of the soil samples was undertaken at the Department of Soil, Crop and Environment Science at the University of the Free State, South Africa. Soil texture was determined in seven fractions using wet sieving for the sand fractions and a hydrometer for the clay and silt fractions. Chemical analyses included organic carbon, and total nitrogen (using dry combustion), pH (water and KCl), electrical resistance, soluble and exchangeable cations, and cation exchange capacity in ammonium acetate, buffered at pH 7. (Exchangeable cations refer to those cations adsorbed to clay and other colloid surfaces that are exchangeable by adding another cation, e.g. ammonium in this case). All analyses were done using standard procedures detailed in The Non-Affiliated Soil Analysis Work Committee [31] and are detailed in [19].

### Rainfall, Meteorological, Climatological Data and Temperatures

Rainfall and other climatological data were collated from two sources: The African Rainfall Climatology 2.0 (ARC2) created by National Oceanic and Atmospheric Administration Climate Predicting Center (NOAA/CPC).

The daily/cumulative rainfall for six selected sites from the Africa Rainfall Climatology (ARC) data, are shown in Figure 2, Daily rainfall for the 2017-2018 (grey lines) season shows peak rainfall is between January and March. Rainfall for 2015-2016 season (blue) across all sites was far below the long mean rainfall (red line).

**Figure 2.**
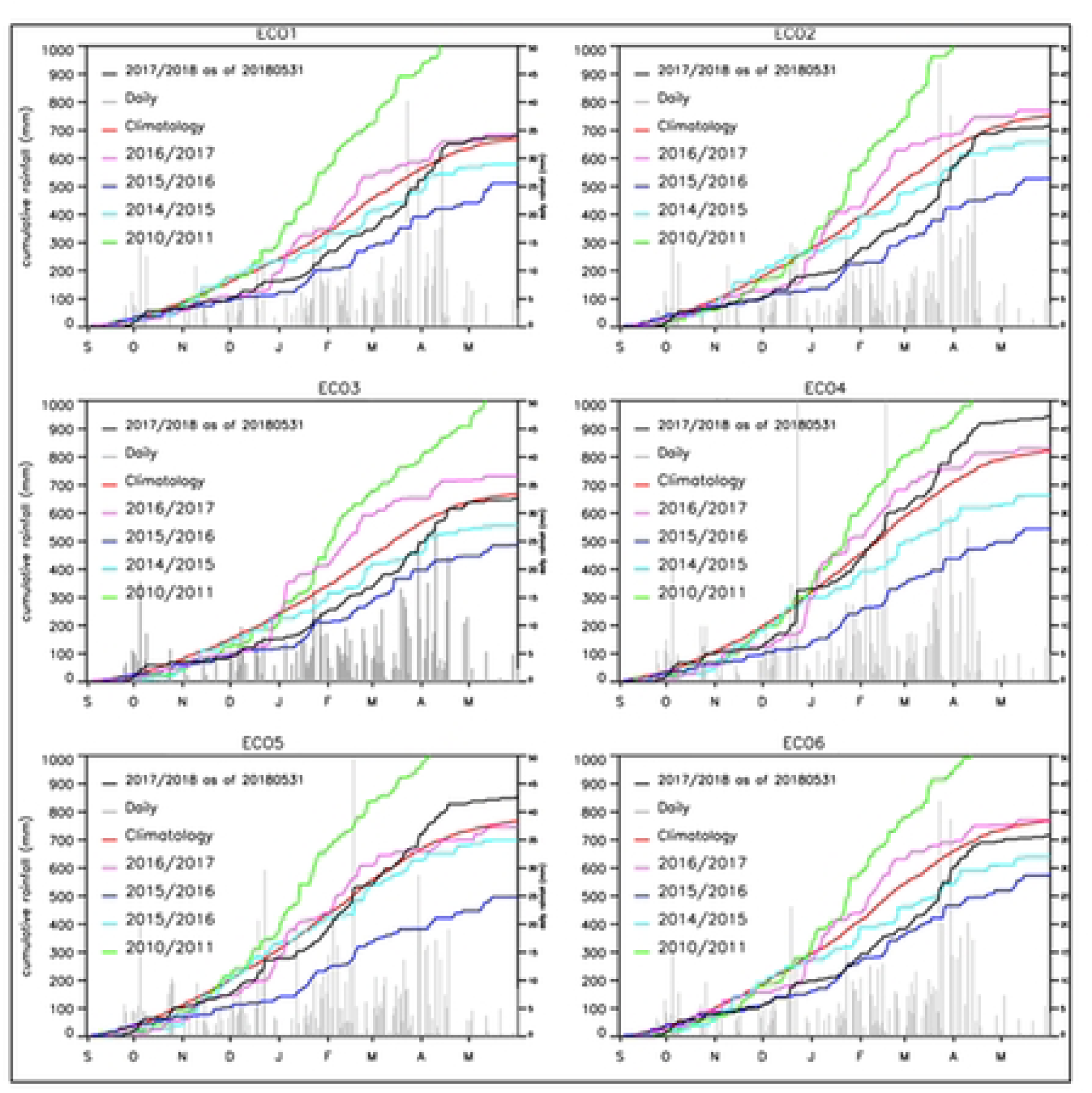
Cumulative rainfall from 2010/11 to 2017/2018, with the highest rainfall for the outbreak in green, at over 1000mm, and the lowest rainfall averaging 500mm, during the drought of 2015/2015, shown in blue. The faint, grey bars show cumulative daily rainfall for each of the months for the full eight years of the climatology data.

### Environmental Sampling

Environmental samples included soil and water. Water temperature was taken using a standard laboratory mercury thermometer (range 0 – 100°C) on the surface of the water and at a 15 cm depth once per site before starting the vegetation transects. Ad hoc, in-sun, ground, spot-temperatures were taken on various substrates using an infrared, hand-held thermometer (Major® MT 691 InfraRed thermometer).

### Vegetation Surveys and Sampling Techniques

Fieldwork was carried out from 2016 to 2018 with initial identification of known plants done in the field and later confirmed in the Geo Potts Herbarium (BLFU) at the University of the Free State. Problematic material was identified at the South African National Biodiversity Institute Herbarium (PRE) in Pretoria. Plant species nomenclature was according to Germishuizen *et al.* (2006), and updated with the 2014 PRECIS database at SANBI, Pretoria. Habitat as well as floristic data was captured using VegCap [32].

The classification of the wetland vegetation was ‘azonal’ as per [33]. The definition of wetlands due to hydrogeomorphic conditions producing redox, anaerobic soils, and difference between wetlands and depressions are discussed in detail in Brand *et al.* (2018) [20].

A combination of two vegetation sampling techniques was used in this study. We used the 1) Braun-Blanquet method (1924, 1964), which was used to conform with the first study from 2014 to 2015 ([20], in combination with 2) a line-point method specifically designed for this survey.

### Braun-Blanquet Survey

Plot sizes varied according to wetland type, location and physical access to sites, and were estimated at 6 x 5 m or 10 x 3 m to total 30 m^2^ to comply with theoretical criteria (Westhoff & Van der Maarel 1980) and established field practice in South Africa [26]. In all sample plots, each species was recorded, all plants counted, and cover was estimated using the modified Braun-Blanquet cover/abundance scale: r, +, 1, 2a, 2b, 3, 4, 5 where the abundance values r (very few individuals), + (few individuals), 1 (abundant), 2a (very abundant, cover < 5%), 2b (5-25%), 3 (25-49%), 4 cover intervals 50-75% and 5 with 75-100%. [27, 34] Whitaker, 1980. Plot size, habitat and floristic data for all 22 sites was the same as the original 15 sites as for Brand *et al.* (2013)[20]. Data was collected using the same methods for the seven new sites.

### Data Processing

Habitat and floristic data were captured using VegCap [35], with the subsequent relevés generated exported as a Cornell Condensed format file (CC!) into Juice version v7.0.28 [28].

Full details of the statistical analysis of the data using TWINSPAN (Two-Way Indicator Species Analysis) algorithm of Hill (1979), and Juice 7.0.28 [28], to produce approximate clustering, six hierarchical levels fidelity using phi coefficient combined with Fisher’s exact test and the modified Braun-Blanquet scale to produce the phytosociological/synoptic table, are provided in [20]. Despite the subjectivity and inaccuracy of the Braun-Blanquet method and the use of non-numerical scores ‘r’ and ‘+’ (r = rare, ‘+’ value is present in low numbers with no cover), which pose computation problems discussed in detail by Podani (2006). The subjectivity of the cover/abundance method as discussed by [36] with group size standardised. The continued use of the Braun-Blanquet method in South Africa is also suggested by [26], and used in a subsequent phytosociological analysis by [20].

### Wetland vegetation classification

Within Juice 7.0.28 [28], the lower threshold values for the diagnostic, constant and dominant species when applying the ‘Analysis of Columns of Synoptic Tables’ function were set to 70, 60 and 50 respectively, while the upper threshold values were set to 80, 70 and 60. Species that exceed the lower threshold were listed while those that exceed the upper threshold were printed in bold in the Juice table. This data was used to compile a full syntaxonomical (S1), with an abbreviated table (Table 1) provided in the main text.

**Table 1.**
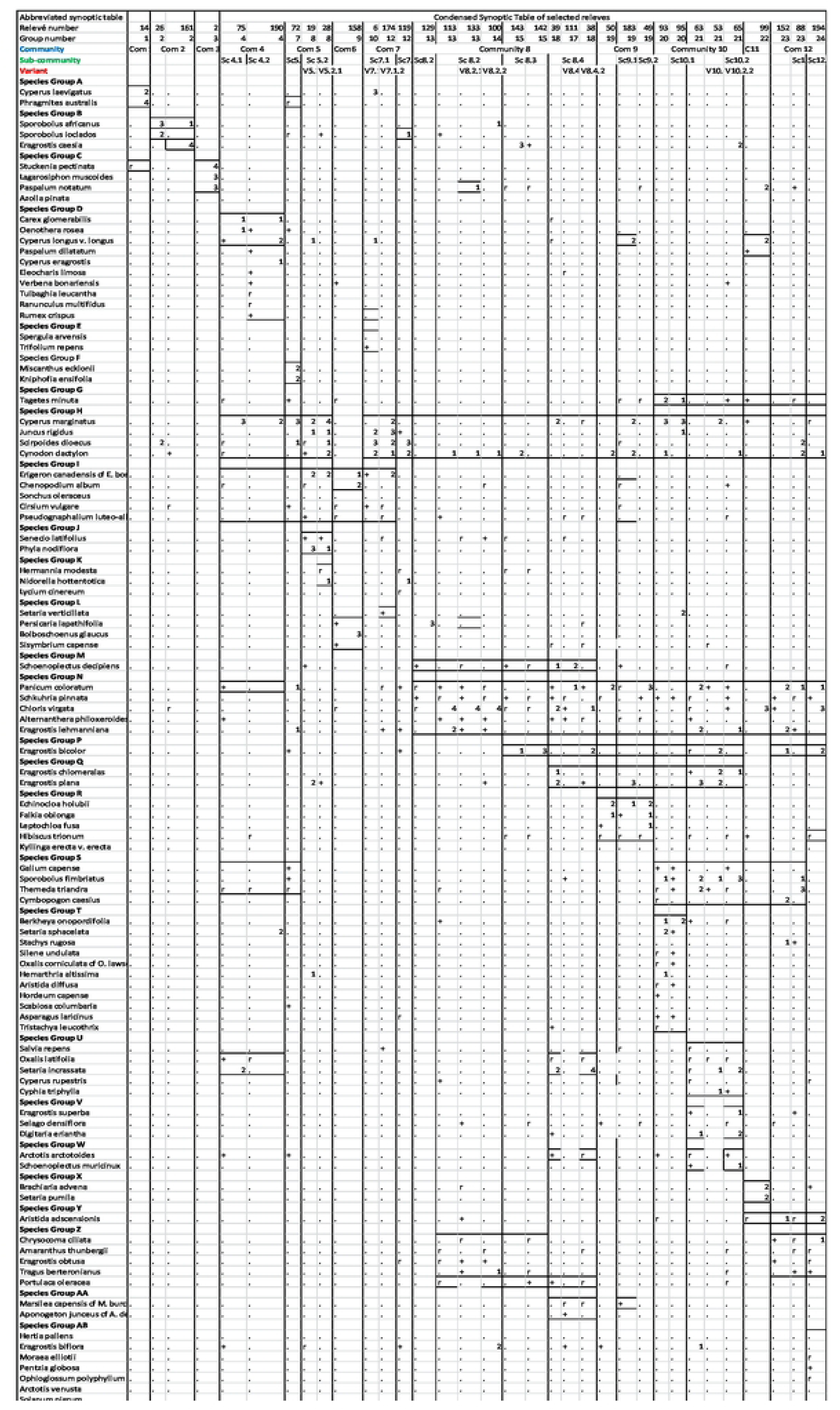
Condensed syntaxonomical table.

### Classification of wetland plant species

This study uses the same wetland-indicator categorization as in [20] and conforms to the standard criteria used to define wetland taxa in [37] Cronk and Fennessy, (2001), Tiner, (1999). These five criteria are discussed in detail in [20].

### Gradient analysis

Groups of similar ecological characteristics were identified and related to environmental gradients. To achieve a normal distribution, the species data were log-transformed during ordination (Legendre & Legendre, 1998), which, in vegetation science, is the ordering, or arranging of plant species or simple units (variants) in a gradient, as plants grow continuously over the landscape. Ordination also summarises information diagrammatically rather than a table, an easier-to-interpret, visual method to display data [27].

A final manipulation of relevé columns and species rows was done in Juice to fine-tune the phytosociological table, which was exported into Excel and refined for presentation by moving rows containing species and adding alphabetic letters to denote species groups (S1 Appendix). The synoptic/syntaxonomical table (S1 Appendix) is the basis of the phytosociological analysis and description. For verification and authentication, a list of all plant species collected by the authors was made (S3 Appendix) and curated by the Geo-Potts Herbarium, University of the Free State [20].

### The line-point (L-P) survey method

To develop our line-point method we integrated two different vegetation methods: line intercept and drop point to obtain vegetation density data and provide quantitative data. We chose to do this following the analysis of the 2014/2015 Braun-Blanquet (B-B) survey [20] as we wanted quantified data to be used in future analyses (Gregory *et al*. In Prep).

The line-point sampling was carried out in conjunction with the B-B plots at the same sites, during the growing seasons, between 2016 and 2018. A total of 201 line-point surveys were conducted, 145 plants were collected, of which 48 had not been collected or identified in the previous study [20]. The combined line-point method is similar, but substantially different with the line intercept method [38]. The line-point method was developed specifically for this study and is a combination of the line intercept method [39] and drop point method of [40]; Figure 3). Six other sampling techniques were compared [41], with the line-point method determined to be the most suitable to assess the vegetation in irregularly shaped, azonal wetlands in the Grassland Biome. The line-point method was the most cost-effective, and allowed for rapid assessments by only one person, and did not need 3 people or cumbersome equipment as illustrated in [40] using the drop point method.

**Figure 3.**
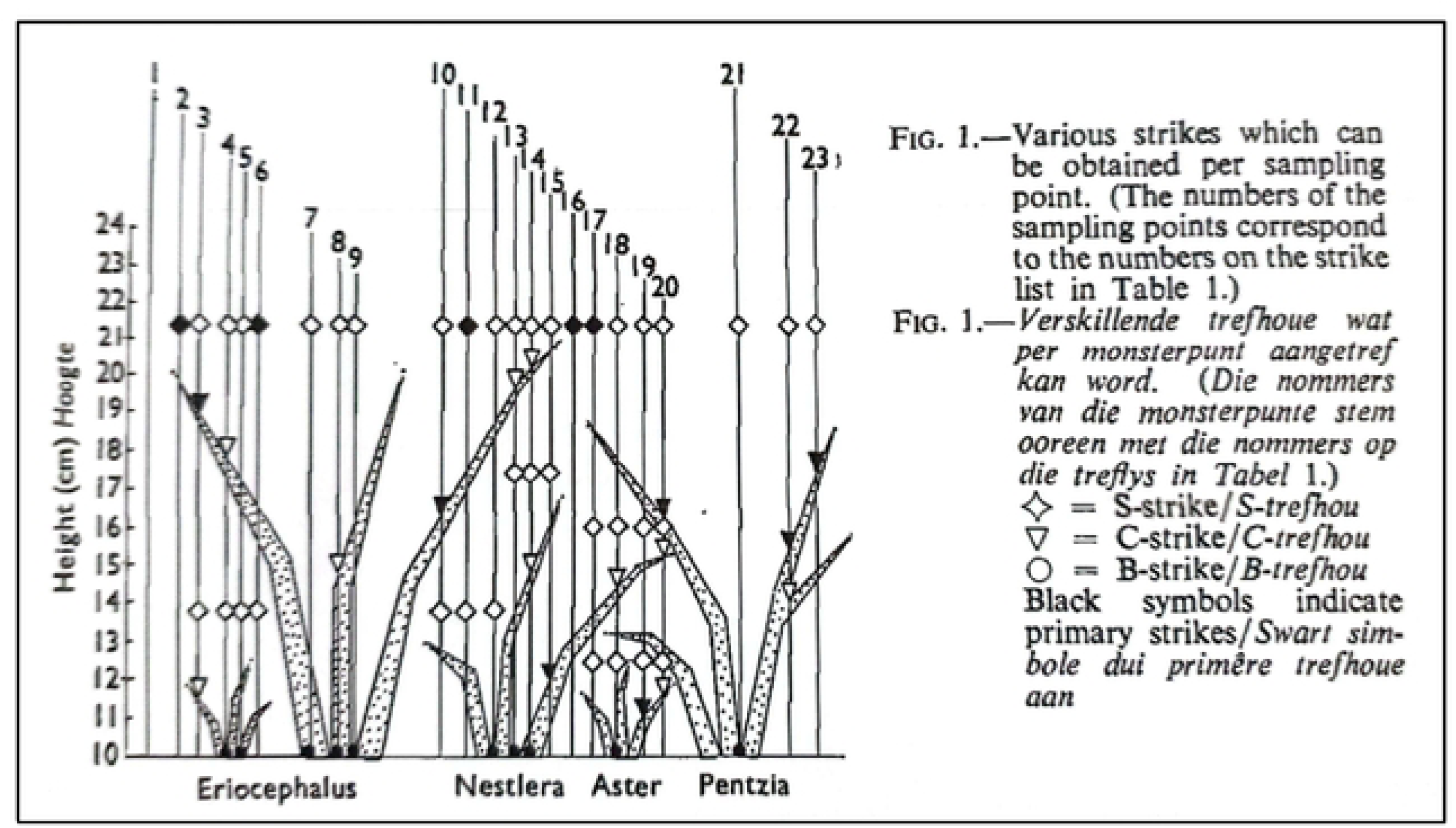
The drop-point method as per Roux 1963, shows how varying strikes measure canopy and height for different plant species. Used according to copyright law expiration timeframe.

The line-point method was designed to measure vegetation density to be used in quantitative analyses (e.g., principal component analysis (PCA). A description of ordination methods is presented in [20]. The traditional line intercept and drop point methods are suitable for open, flat grasslands, where the linear, grid, straight-line layout is possible.

The object of conducting the line-point survey was to ascertain the relationship between the height and density of vegetation, and adult mosquito numbers per site. The taller, or higher the vegetation, the greater the canopy spread, and the more dense the vegetation, the more strike-density per height class were obtained. The wetland areas of interest for the line-point survey consisted of the vegetation found between the lowest to highest areas flooded during maximum inundation (Figure 4).

**Figure 4.**
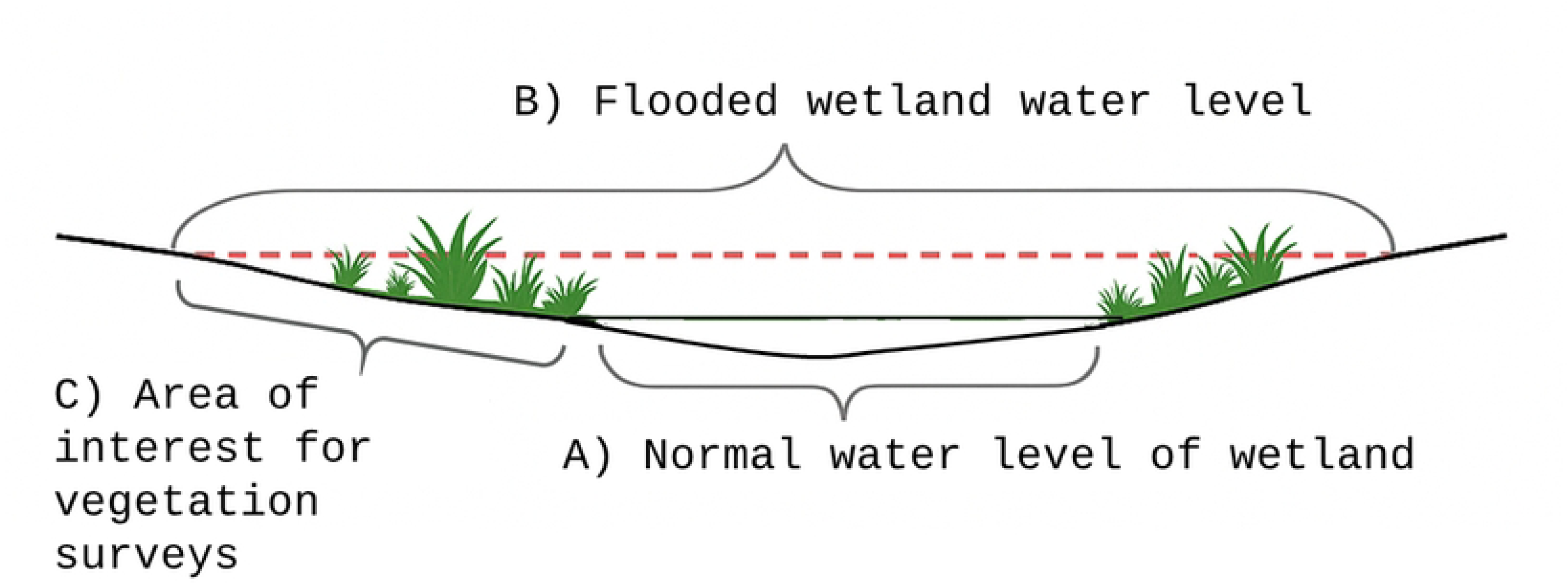
Idealized wetland conditions needed by *Aedes* mosquito species and forms the habitat sampled for the line-point vegetation survey.

The floodwater levels were defined by the life-cycle conditions required by specific *Aedes* mosquito species for laying eggs and for larval and pupal development (Figure 4).

### Theoretical and practical considerations for the line-point method

The azonal wetland areas of interest sampled for the surveys, consisted of vegetation embedded in the Grassland Biome. These wetlands have irregular shapes, with topography of varying slope, with vegetation growing in a variety of very different wetland/stream/pan [20]. To account for these different field conditions, and to still provide the same statistically meaningful data, six layouts were designed and used to sample the vegetation as illustrated in Figures 5 to 7. Figures 7b and 7c, show the L-P method in action. Full descriptions of the various line-point sampling techniques are provided in Appendix D/ Supplement 4.

**Figure 5.**
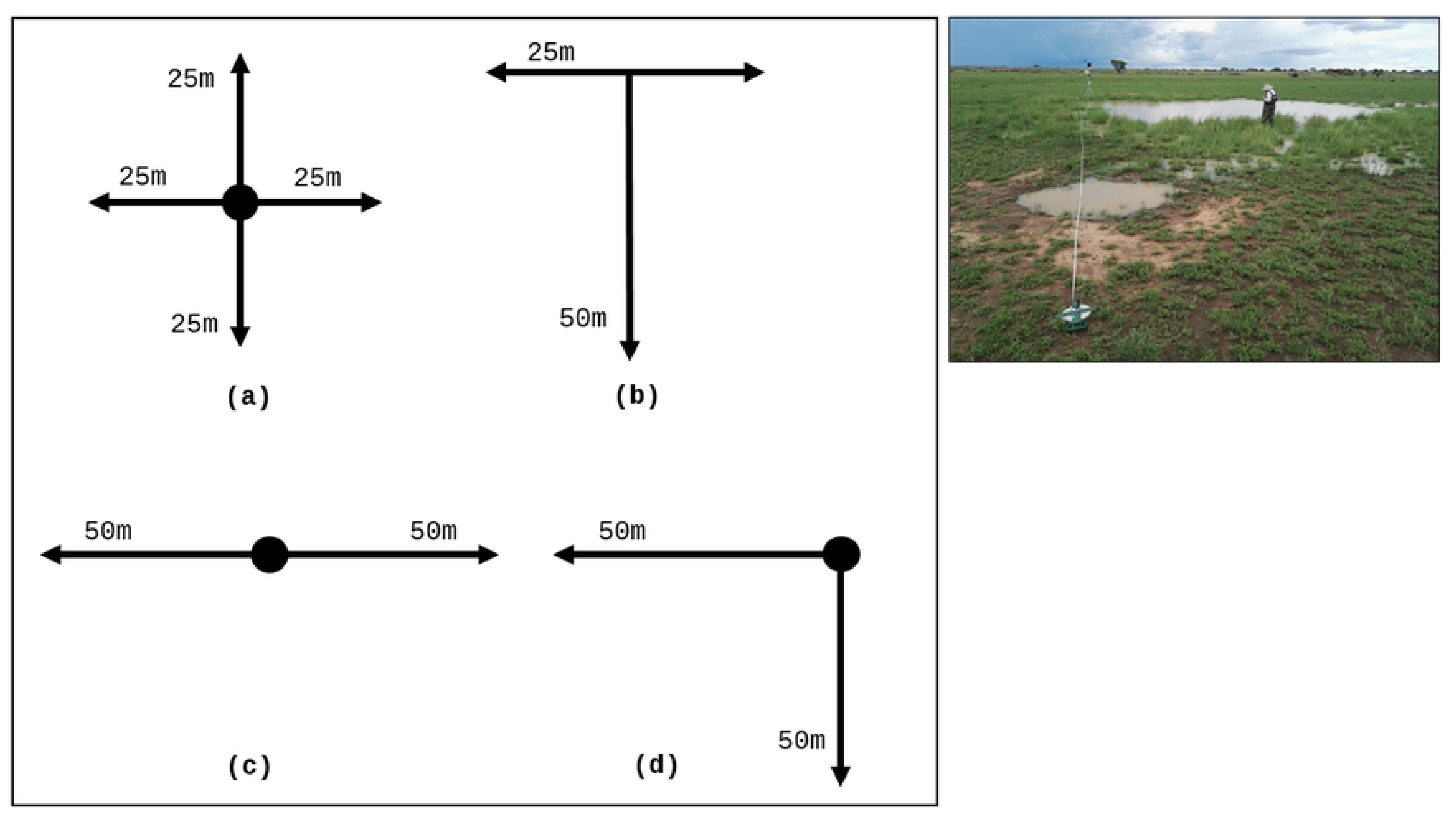
Four possible layouts as determined by theoretical considerations on flat terrain, showing cross (a), linear (b), straight (c), and right-angled (d) transects. The photograph (Figure 5e), shows a 50 m, straight-line transect at site p015. The wetland is located in a large, flat pan, with low, sparse plant cover.

**Figure 6.**
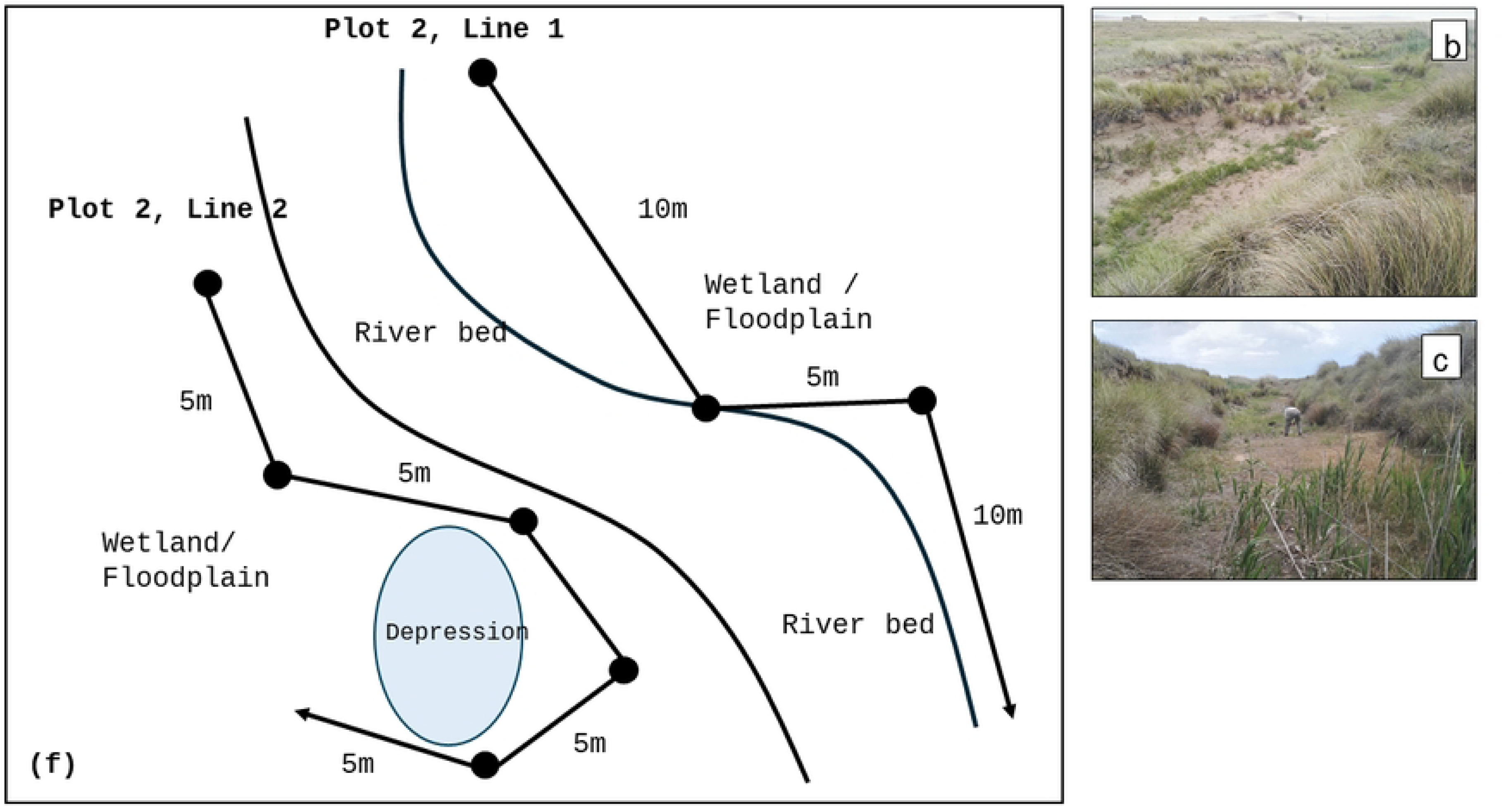
The zig-zag layout for curving riverine or palustrine sites (a), is demarked by the white string, in the wetland shown in the photograph (b, c), at site 131s, Free State, South Africa.

**Figure 7.**
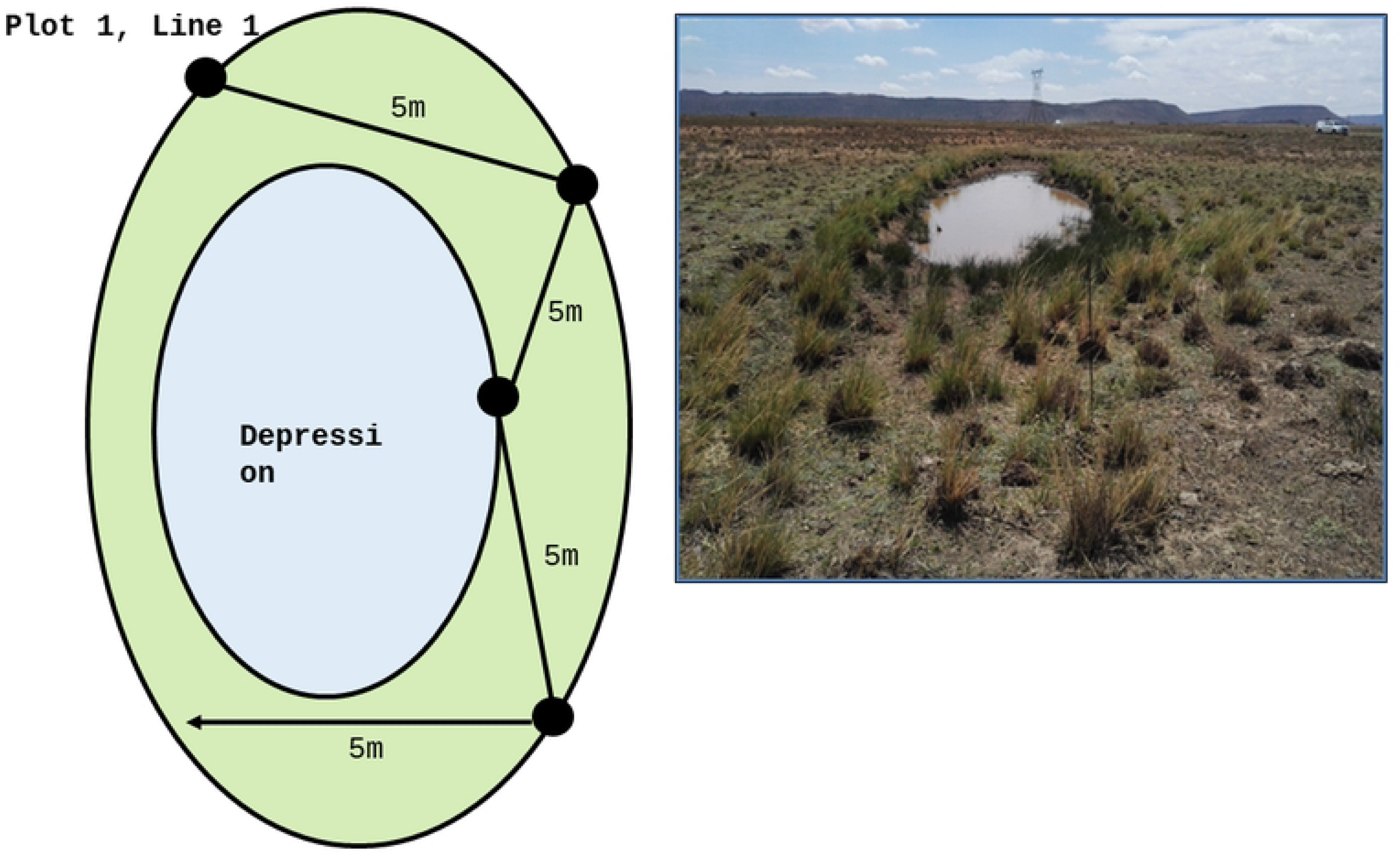
The illustration shows the modified line-point sampling methods used for pans, dams or oxbows. The photograph (b) is of an oval wetland, typical, heavily overgrazed Free State site, which had high sheep mortality during the 2010/11 RFV epidemic. The remaining vegetation show consists of almost impalatable wetland vegetation, *Juncus rigidus*, and sedges, *Scirpoides dioecus*, after all palatable grasses have been eaten.

The L-P method to measure vegetation density of incised, seasonally dry streams, with wetland flooded pans are illustrated in Figure 5a, with photographs documenting the white string demarking the transects, shown in Figure 5b and 5c.

The zigzag transect-pattern is used in irregular, incised, palustrine wetland with dense vegetation. The bamboo-like grass, *Phragmites australis* is in the foreground, dense stands of the sedge, *Scirpoides dioecus* grow on the sides, and the tall, bunch grass, *Miscanthus ecklonii*, grows on the top of the wetland trench.

For typical ovoid-shaped wetlands, we used an m-shaped transect (Figure 6). These ovoid-shaped oxbow wetlands are formed as part of the Palaeo-Kimberley river system [42], and are typical of shallow upland depressions, only inundated during the rainy season, found throughout the study area, and more broadly throughout the interior of South Africa.

Free State oxbows are the result of a mature riverine landscape that, since the Triassic era, has only had the geological process of weathering resulting in the erosional levelling of the landscape, allowing for stream-meandering which produces oxbows [43] (Goudie, 1991).

### Mosquito sampling

Four CDC or two modified Shannon traps were placed adjacent to each pan and baited with dry ice (CO_2_) to trap mosquitoes between dusk and dawn. Mosquitoes were killed by freezing and transferred on dry ice to the National Institute for Communicable Diseases of the National Health Laboratory Service (NICD-NHLS), where they were stored at -70°C. Mosquitoes were identified morphologically using taxonomic keys [23, 44, 45] and sorted by species for storage in cryotubes at -70°C. Relative mosquito abundance was approximated as trap rate, i.e., the total number of mosquitoes per trap-night, and plotted against average (arithmetic mean) monthly rainfall across all sampling localities. Sampling locality data were plotted as quadrants of the approximately 400,000km^2^ study area.

## Findings/Results

As per [20], the skewness and kurtosis calculations performed with PC-ORD v5.0 revealed the non-unimodal distribution of the species data (also confirmed by the disjunct nature of the dataset as indicated by the Detrended Correspondence Analysis (DCA) eigenvalue of one for the first axis. The Detrended Correspondence Analysis (DCA) produced the Ordination Diagramme (Figure 8), and is a reliable statistical method in multivariate data analysis, to retain as much of the variation as possible, and which shows the relationship of the identified plant communities with the environmental variables.

**Figure 8.**
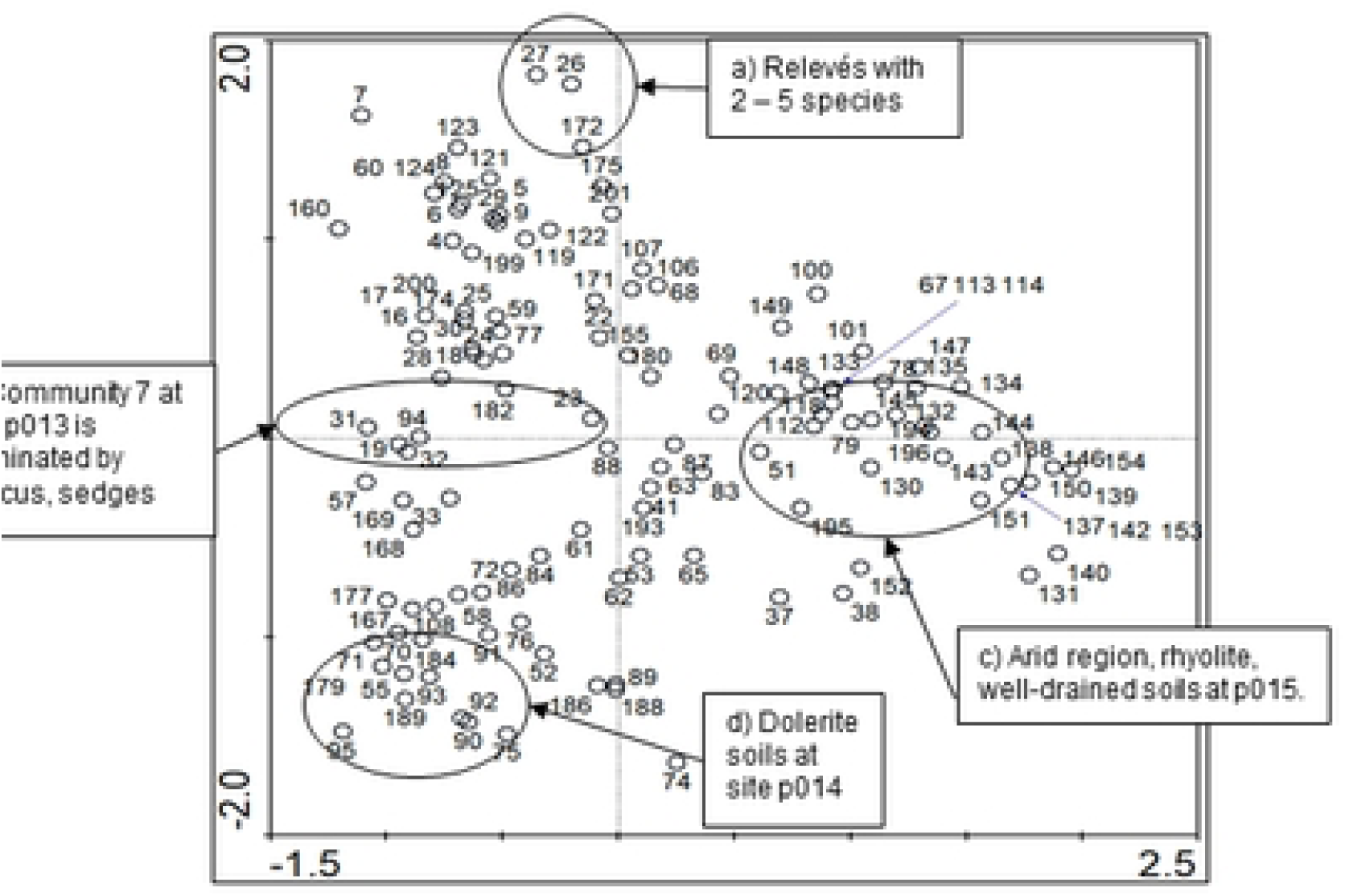
The ordination diagramme is the third iteration after removing tightly overlapping relevés to provide a clearer indication of local environmental conditions; a) shows clusters with only 2 – 5 relevés, possibly inundated wetlands or depauperate communities due to overgrazing. b) These relevés had dominant and diagnostic wetland species of sedges and *Juncus* and represent sites of the highest RVF mortalities. c) were relevés on the rhyolite volcanic soils in the arid most westerly study sites situated in the Savanna Biome. This region had the lowest rainfall, with different soils from the rest of the study sites located on the Karoo-Supergroup geology. d) Relevés occurred on dolerite outcrops with different dominant and diagnostic species, with no or few RVF mortalities.

### Ecological and environmental variables

The ordination diagramme (Figure 8), illustrates the gradients of ecological and environmental conditions.

The ordination analysis (Figure 8), indicates there is no clear single or set of biotic and abiotic ecological conditions that acts as main driver/s to determine the most significant vegetation/soil conditions responsible for *Aedes* mass breeding as represented by the reporting of historical RVF mortalities.

### Soils

The dominant reference soil groups were Kastanozem and Calcisol, with Calcic, Haplic, and Hypocalcic; and either Clayic or Loamic as supplementary qualifiers [46]. For the top soils, the organic carbon ranged from 5.1 m kg^-1^ to 76.7 g kg^-1^, while total nitrogen ranged from 0.30 g kg^-1^ to 4.74 g kg^-1^. Soils were rather alkaline, with pH_water_ of the topsoil ranging between 5.44 and 10.04, whilst also being predominantly saline as evidenced by the resistance that varied between 12 and 7 400 Ω. Exchangeable calcium ranged from 6.50 cmol_c_ kg^-1^ to 61.1 cmol_c_ kg^-1^, exchangeable magnesium from 1.18 cmol_c_ kg^-1^ to 23.8 cmol_c_ kg^-1^, exchangeable potassium were very low and ranged from 0.13 cmol_c_ kg^-1^ to 6.2 cmol_c_ kg^-1^, while exchangeable sodium ranged 0.12 cmol_c_ kg^-1^ to 26.1 cmol_c_ kg^-1^. The cation exchange capacity (CEC) also varied widely from 3.83 cmol_c_ kg^-1^ to 41.7 cmol_c_ kg^-1^. Detailed descriptions and analytical results of a subset of sites are given by [19].

### Wetland-Types

A categorization of wetland-types was presented in [20], and consisted of five categories of wetlands; 1. Endorheic salt pans, 2. Non-saline depressions, 3. Palustrine wetlands, 4. Riparian wetlands, 5. Anthropogenic Wetlands. No new categories were found for this study, with all seven new sites surveyed falling with categories 1 to 4. Of the eight new sites, two of them do have earth-walled dams blocking existing rivers and streams, and, arguably, could be defined as ‘anthropogenic’, however, they were not, the distinct category 5, Anthropogenic Wetlands, resulting from irrigation canal-type wetlands, found in western Free State, which are there exclusively due to the irrigation canal.

### Floristic analysis

#### Widespread wetland plant community species

Numerous species were found to have widespread distribution with some being common through the wetland study sites. The majority were grouped in associations, but some were single species e.g., *Phragmities australis,* the largest Obligate Wetland graminoid macrophyte, (Species Group A), found with low cover/abundance scattered throughout relevés comprising communities 1 to sub-community 7.1.

Species Group H, was comprised of *Cyperus marginatus, Juncus rigidus, Scirpoides dioecus, Cynodon dactylon*. It was a broad, but scattered group of constant species found through all communities except Community 3.

Other, less common but widespread taxa, formed Species Group I: *Erigeron canadensis* c.f. *E. bonariensis, Chenopodium album, Sonchus oleraceus, Cirsium vulgare* and *Pseudognaphalium luteo-album*.

Species Group N was composed of: *Panicum coloratum, Schkuhria pinnata, Chloris virgate, Alternanthera philoxeroides* and *Eragrostis lehmanniana* are common to many communities excluding communities 1 to 3.

The aromatic, herbaceous, perennial, Lamiaceae forb, *Salvia repens*, Species Group U, was scattered throughout many relevés, but with values of only ‘r’ for rare, it forms a distinct community with other members of species group U, as shown in sub-community 10.2.

#### Phytosociological analysis and syntaxonomic ranks

The vegetation survey consisted of 201 relevés. We identified 12 vegetation communities as presented in the full synoptic/syntaxonomical table (S1). There were 18 sub-communities and 10 variants. For brevity, only community associations are described in the main text. The full phytosociological description of all 40 vegetation (sub)communities/variants is presented in Appendix C/Supplementary S3. Five relevés located at the extreme right of the synoptic table do not represent any significant vegetation associations, regarded as ‘noise’, and were not included in the description. For ease of reference a condensed version of the synoptic table showing the syntaxonomic ranks is presented in Table 1 (condensed syntaxonomical table). A total of 194 plant species were identified, with species names, authors and botanical voucher numbers identified and presented in Appendix B/Supplementary S2. Recent changes in vegetation science have changed the previous use of phytosociology to use syntaxonomy and show syntaxonomical rank on vegetation tables [26]. Syntaxonomy and syntaxonomical rank refers to the concept of using a hierarchical system of ranking vegetation associations from highest to lowest; class or order for Europe [47], and in South Africa, community, sub-community and variant [26].

In phytosociological studies, it is recommended that formal syntaxonomic descriptions are given (Brown *et al*., 2013), which also allows for an abbreviated version of a syntaxonomical/phytosociological table to be presented in the main body of the text, with the complete table show in an appendix as supporting information. The wetland vegetation is embedded in the matrix of the surrounding Grassland biome, and, as this study had a limited coverage, is more localized and has smaller data sets within the 40 000 km^2^.

#### Naming the plant communities

All 40 vegetation associations are named, showing diagnostics, co-diagnostic, dominant, co-dominant and common species. The plant community names and physiognomic descriptors are presented in full in Supplementary Material 3, Appendix C. The physiognomic name, e.g., inundated wetland, wetland/emergent grassland depression, follows criteria in [32, 48] South African Department of Water Affairs and Forestry, (2003), Brand *et al*., (2013).

#### Describing the plant communities

The results of the classification produced 12 communities, 18 sub-communities and 10 variants, which are described using the diagnostic species first, followed by the dominant species (Brown *et al*., 2016). Some species are found throughout several vegetation associations, and form wide-spread clusters which may be used as dominant or diagnostic species. Full and detailed phytosociological descriptions of the remaining 18 sub-communities, and 10 variants are provided in (Appendix C; S3).

Overall, the syntaxonomical table (Table 1; and S1) showed a clustering for wetland species to the left in the condensed table and Appendix 1, (communities 1, 2 and 4), with community 3 (SG C), composed of submerged aquatic species (*Lagarosiphon muscoides*), an Obligate Wetland species (OBL). Community 8 (SG M and N) was the largest community, composed mostly of Facultative (FAC), Facultative Upland (FACU), with a few scatterings of Facultative Wetland (FACW) plants showing limited wetland syntaxa. The degree to which plant species associate with wetlands and their wetland-indicator status is detailed in Cronk & Fennessy (2001) and Tiner (1999), and categories wetland plants; (OBL, FACW, FAC, FACU) and species, based on estimated probability of finding plants in these locations. Details are discussed in [20].

Communities 11 and 12 were composed of mostly Upland and Facultative Upland species (Appendix A: S1).

#### Line-point analysis and results

A total of 201 line-point transects were sampled. High density vegetation was exemplified by the megagraminoid *Miscanthus capensis*, a tall, densely tufted grass which can reach heights of 2.7 m, and has a canopy spread of 0.8 to 1.4 m (Figure 9; ad hoc field measurements) [49]. Low density vegetation was exemplified by the dense, wiry sedge, *Scirpoides dioecious* can average 0.8 to 1.2 m in height, and spread 0.4 to 0.8 m (Brand *et al*., 2016, ad hoc field measurements). Both graminoids form dense stands which provide protected, cool, moist habitat for adult mosquitoes. Abbreviations of plant genera are given in Appendix A and C.

**Figure 9.**
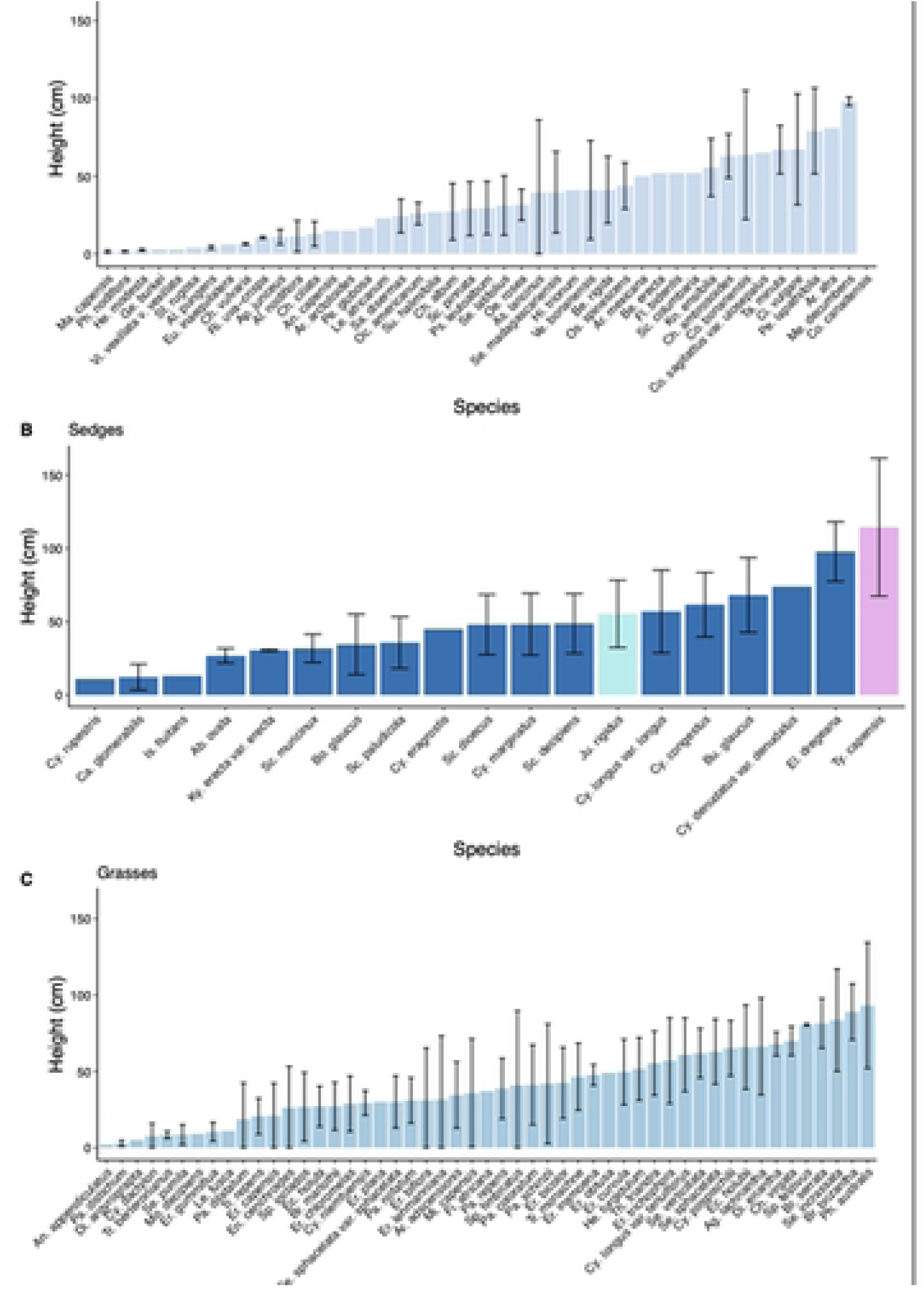
Line-point surveys results showing the mean height for species within three discrete vegetation classes of a) forbs, b) sedges and c) grasses. The black bars represent average variation in height of each plant species. If no black bar is on the plot, then only one identification was made of that species. In the sedge height class, for convenience, two non-sedges are included, these are *Juncus rigidus* (light blue bar), and *Typhae capensis* (Typhaceae; Ty. capensis; pink bar), which form the tallest vegetation community (1), even though Phragmites australis is the tallest graminoid with a range varying 50 to 140 cm.

Strike density ranged from 0-0.36 strikes per 10cm and strike height ranged from 0.25-20 cm during the wettest months of the growing season, January to March (Figure 10).

**Figures 10.**
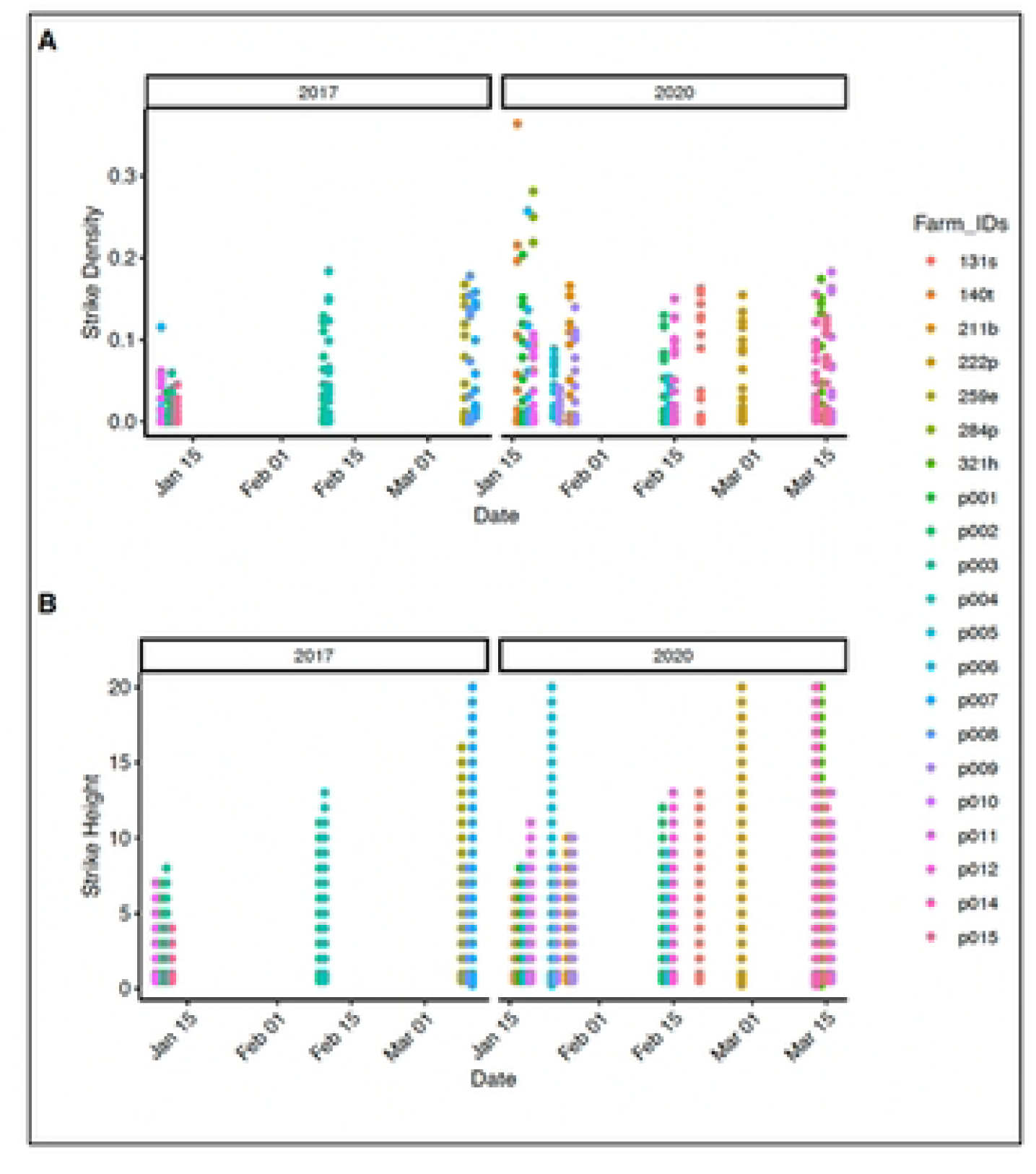
Mean strike height and strike density per site in 2017 to 2020 after the wetland vegetation has recovered from the 2015/16 drought.

#### Mosquito sampling and site collection

A total of 111,312 adult female mosquitoes were sampled and identified for this period, yielding 57,583 floodwater *Aedes* spp. and 37,770 *Culex* spp. amongst species of lesser interest.

Relative mosquito abundance, measured as trap rate (number of mosquitoes per trap-night), varied with the amount of local rainfall at each study site. The relative mosquito abundance increased from 2014 to 2018 (Figures 11 a, b, c, d). At lower rainfall levels, floodwater species such as the RVFV vectors emerge only in small numbers and, possibly, only at a few sites at any one time. A detailed assessment of the mosquito population richness, abundance and turnover rates is in prep (Gregory *et al.*, In prep).

**Figure 11.**
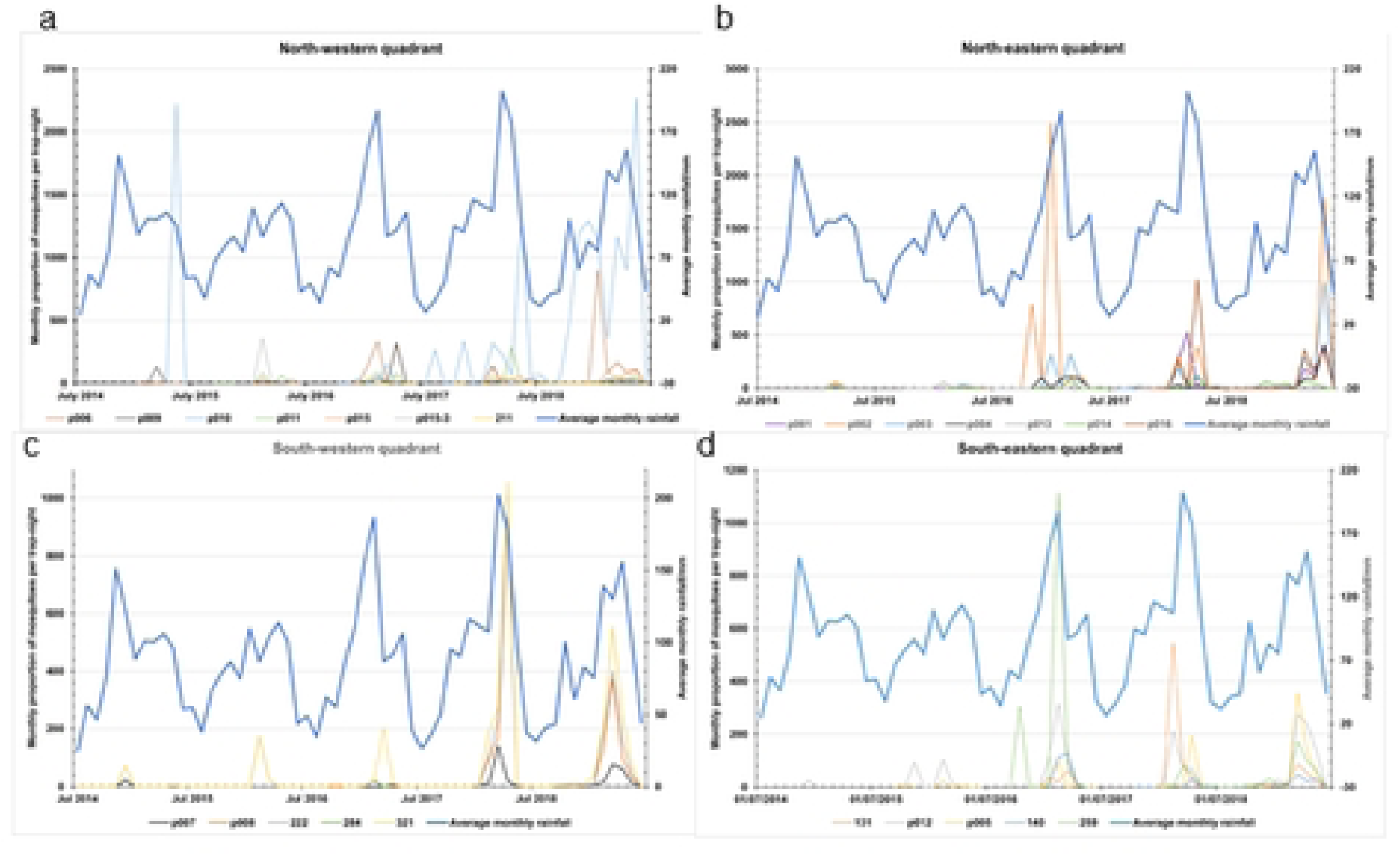
Four quadrants of the study area showing total mosquito numbers per rainfall plots.

As the rainfall levels increase from drier to wetter years, the mosquito relative abundances increased.

#### Temperature and moisture associated with wetland vegetation

While assessing the vegetation for each the study sites, *ad hoc* temperatures were taken with a hand-held thermometer using an infrared beam. Temperatures were obtained on the top of plants, on the sides, on the bare earth, on dead plant material and under the plant canopy. Results in Figure 12 show a maximum range from 71 - 80°C (179°F), in full sun, on top of plants, on dead grass-tufts and on bare earth. Temperatures varied on the side of the tufts from 38 - 45°C, and under the plant canopy, in the growth-point center of the plant, from 25 - 32°C.

**Figure 12.**
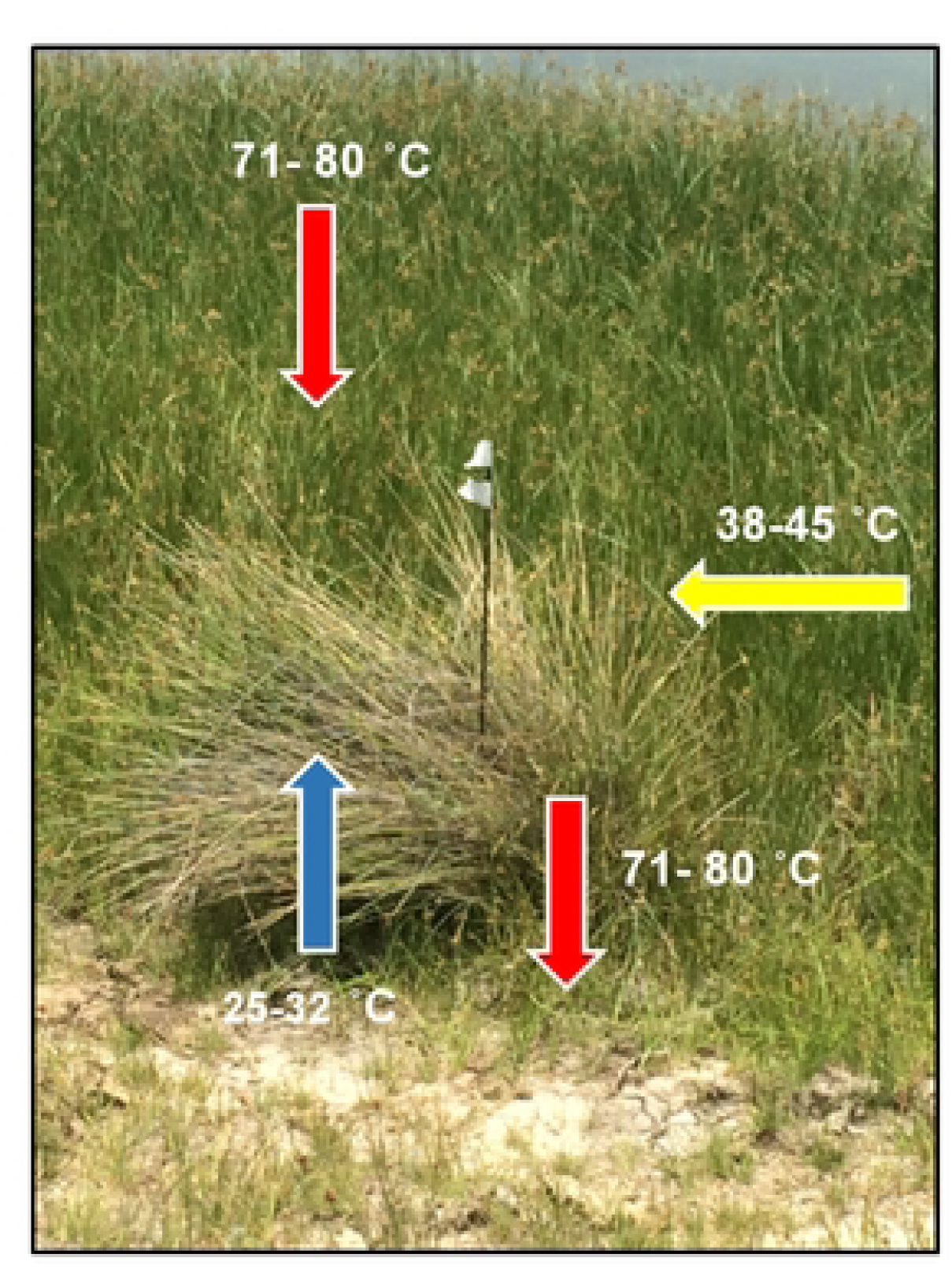
Temperatures measured on the sedge *Scirpoides dioecus* and substrate in full sun at noon show temperature from 25°C to 80°C, a range of 55°C. The black steel dropper with white tape, marks a line-point transect starting-point.

#### Combining wetland categories, vegetation, soils and floodwater Aedes

We described the category, wetland, dominant vegetation, geology and soil and predominant floodwater mosquito vectors for all sites (Table 2). For sites that had vegetation characterization in 2014/15 [20] and 2017/18, we also compared the results of the two surveys (Table 2). We documented a change in community structure in all the 15 previously surveyed sites, among species composition and associated dominant and diagnostic species. Over-grazing and erosion that resulted from the drought in 2015/6, may be one possible explanation for the change in plant structure and associations, between the two studies.

**Table 2.** Combining sites with high sheep mortality, wetland category, dominant vegetation, *Aedes* mosquitos, geology and soils, Free State, South Africa.

| Farm identification number | Cate - gory | Wetland category | Dominant vegetation | Geology, soils, notes | Predominant floodwater mosquito species |
| --- | --- | --- | --- | --- | --- |
| p015 | 1 | Depressions, endorheic non-saline pans. | grass/<br>invasives; no OBL species | Andesite lava & modern, red Kalahari aeolian soils | *Ae. mcintoshi*, *Ae. caballus*, *Ae. juppi*, *An. squamosus/cydippis* |
| p013 | 1 & 2 | Upland depressions, possibly palaeo-river. | Sedges<br>*/Juncus* | Endorheic salt pan. Calcrete outcrops. Ecce, Beaufort shale. | *Ae. caballus*, *Ae. juppi*, *An. squamosus/cydippis* |
| p004 | 1 & 2 | Permanent wetland embedded in salt pan. | Sedges<br>*/Juncus* | Calcrete. | *Ae. mcintoshi*, *Ae. caballus*, *Ae. harrisoni*, *Ae. juppi*, *An. squamosus/cydippis* |
| p011 | 1 & 2 | Shallow depression, open salt pan. | Sedges<br>*/Juncus* | Saline pan, Calcrete. | *Ae. mcintoshi*, *Ae. caballus*, *Ae. juppi*, *An. squamosus/cydippis* |
| p001 | 2 | Upland depressions, possibly palaeo-river. | Sedges/grass | Ecce, Beaufort shales | *Ae. mcintoshi*, *Ae. caballus*, *Ae. juppi*; *An. squamosus/cydippis* |
| p002 | 2 | Depression, palaeo-rivers, seasonally inundated | Grass/sedge/<br>*Juncus* | Karoo sediments, Ecce, Beaufort group grey shales | *Ae. mcintoshi*, *Ae. caballus*, *Ae. juppi*, *An. squamosus/cydippis* |
| p008 | 2 & 3 | Palustrine wetlands. Depressions | Sedges/grass | Clay soils | *Ae. caballus*, *Ae. juppi*, *An. squamosus/cydippis* |
| p005 | 2 & 3 | Palustrine wetlands. Depressions | *Juncus*/<br>*Phragmites* | Calcrete, Hornfels with dolerite fragments | *Ae. caballus*, *Ae. juppi*, *An. squamosus/cydippis* |
| p006 | 2 & 3 | Depression, palaeo-rivers, and palustrine wetland | Sedges/grass | Calcrete, shale, red aeolian sandy soil. Gray Beaufort shales | *Ae. mcintoshi*, *Ae. caballus*, *Ae. juppi*, *An. squamosus/cydippis* |
| p003 | 3 | Palustrine seasonally inundated | Sedges | Karoo Super-group. Ecca, Beaufort Sandstones, grey shale | *Ae. mcintoshi*, *Ae. caballus*, *Ae. juppi*, *An. squamosus/cydippis* |
| p009 | 3 | Artesian spring-fed wetlands | Sedges/<br>Juncus | Calcrete 3-4 m thick | *Ae. mcintoshi*, *Ae. caballus*, *Ae. harrisoni*, *Ae. juppi*, *An. squamosus/cydippis* |
| p014 | 3 | Palustrine wetlands | Sedges | Extensive Dolerite sills | *Ae. caballus*, *Ae. harrisoni*, *Ae. juppi*, *An. azevedoi*, *An. squamosus/cydippis* |
| p012 | 3 & 4 | Palustrine wetlands. Riverine wetlands | Sedge/grass/<br>Juncus | Clay soils from | *Ae. dentatus*, *Ae. hirsutus*, *Ae. luridus*, *Ae. mcintoshi*, *Ae. unidentatus*, *Ae. caballus*, *Ae. juppi*, *An. squamosus/cydippis* |
| p010 | 5 | Anthropogenic floodplain, irrigation dam overflow | Sedges/grass | White, salt-leached soils | *Ae. caballus*, *Ae. juppi*, *An. squamosus/cydippis*, *Cq. microannulata* |
| p007 | 5 | Anthropogenic wetland made by pivot irrigating | Sedge/grass | Endorheic saline pan & upland depression, Ecca & Beaufort shale | *Ae. caballus*, *Ae. juppi*, *An. squamosus/cydippis* |
| *Eight new sites added in 2016:* |  |  |  |  |  |
| 259e | 2 | Palustrine wetlands, possibly palaeo-river. Upland depression | Grasses dominant, limited Juncus | Karoo-Supergroup, Ecca, Beaufort shale | *Ae. luridus*, *Ae. mcintoshi*, *Ae. caballus*, *Ae. juppi*, *An. squamosus/cydippis* |
| 140t | 2 | Palustrine wetlands, possibly | Grasses dominant, limited Sedge | Karoo-Supergroup, | *Ae. mcintoshi*, *Ae. caballus*, *Ae. juppi*, *An. squamosus/cydippis* |
|  |  | palaeo-river.<br>Upland<br>depression |  | Ecce & Beaufort<br>shales |  |
| 131s | 2 | Palustrine<br>wetlands.<br>Depressions | Grasses and<br>Sedge,<br>*Scirpoides*<br>*dioecus* | Karoo-Supergroup,<br>Ecce & Beaufort<br>shale | *Ae. luridus, Ae.*<br>*mcintoshii, Ae.*<br>*unidentatus, Ae.*<br>*caballus, Ae. harrisoni,*<br>*Ae. juppi, An.*<br>*squamosus/cydippis* |
| *321h | 2 | Palustrine<br>wetlands.<br>Depressions | Grasses<br>dominant;<br>*Phragmites*;<br>*Juncus* | Karoo-Supergroup,<br>Ecce & Beaufort<br>shale | n/a |
| 284p | 5 | Anthropogeni<br>c dam,<br>shallow<br>depression | Grasses only | Karoo-Supergroup,<br>Ecce & Beaufort<br>shale | None |
| 211b | 2 & 3 | Shallow<br>upland<br>depression,<br>palustrine<br>wetland | Grasses only | Karoo-Supergroup,<br>Ecce & Beaufort<br>shale | *Ae. caballus, Ae. Juppi* |
| 222p | 3 & 4 | Palustrine<br>wetlands.<br>Riverine<br>wetlands | Grasses<br>dominant;<br>*Phragmites*,<br>*Miscanthus*<br>*sp.*,<br>Juncus/Sedge | Karoo-Supergroup,<br>Ecce, Beaufort<br>shales | *Ae. caballus, Ae. juppi,*<br>*An.*<br>*squamosus/cydippis* |

## Discussion

This study provides a detailed description of vegetation, soil, climate and vector data from a region that is susceptible to large outbreaks and where small and localized RVF epizootics occur regularly [50]. Our findings help to understand the complex ecosystem that supports floodwater *Aedes* and consequently RVFV maintenance. We have shown the harsh conditions at which the mosquito eggs must survive (vegetation and soil temperature up to 80°C) are likely mitigated by vegetation communities that provide shelter for survival until the next flooding period. Vegetation has an important influence on varying absorption rates of infrared radiation (Bauer *et al.,* 2021) and provide resting sites for mosquitoes during the heat and aridity of the day. We have also shown how these conditions change over a relatively short period of time. We suggest that more detailed ecological analyses are necessary to understand the mechanisms of RVFV survival in harsh conditions and identify indicators of change that may result in an outbreak of RVF.

### Soils and vegetation

Detailed soil analyses done by Verster *et al*. 2020 [19] at 15 sites and [30] Gqalaqha (2020), found considerable statistical variability of soil mineralogy and microbiology between the sites that reported RFV mortality and sites that had no mortality.

The initial study of [20], used visual field assessment to identify high clay-content soils. This was not corroborated by the detailed laboratory findings of Verster *et al*., (2020) [19], which found a lower medium sand content. Medium sand and not clay content, may be a better predictor of wetlands as breeding sites for *Aedes*. Sandy soils allow for better water drainage after heavy rains, rather than high clay-content, water-retaining soils, and thus provide a more suitable *Aedes* breeding habitat. Additionally, *Aedes* mosquito larvae require sandy soils to clean their mouth parts [22] indicating better water retention in the pan substrate.

All five wetlands categorized from the previous study [20] were freshwater wetlands. However, there are two main divisions; natural, freshwater wetlands and anthropogenic freshwater wetlands. The importance may lie in the difference in soils [19, 30] and the habitat necessary for the eggs to be laid and the larval stages to develop (Dale & Knight, 2008). This will also include the ephemeral, palustrine and shallow depression sites which dry up during the non-rainy season. The floodwater *Aedes* are obligate pan-breeding species, unlike the *Culex* spp., which only utilise the pan wetland opportunistically and typically breed in more permanent waterbodies (Alan Kemp pers. comm.).

The distinct plant associations identified at site p015 (Holpan/Graspan National Park, [20], occurring on soils derived from rhyolite, were not clearly identified in this study. However, there were more invasive species identified in this study. It may be that these species are interspersed and not show as discrete communities in the syntaxonomical table (Table 1; Supplementary Material, Appendix A) as in the previous study of [20].

Additionally, the 8 new sites where not selected for wetlands, but rather for serum studies on sheep. These sites were overgrazed and formed significant invasive, non-native, plant-species communities.

The results shown in Table 2 and in Figure 8 (ordination diagramme), imply a high rate of tolerance by the vectors for the variety of wetland categories, soil and vegetation present in the study area. Compared with the situation in Kenya, with significant outbreaks happening in the moist western uplands of Nakuru County (Lake Naivasha at an average of 1896m asl) as well as in the arid eastern Garissa County/Tana County below 400m above sea level (asl), it seems the main drivers of RVFV ecology in South Africa are endorheic pans, occasionally sufficient rainfall and a rich supply of bloodmeals for the vectors [21]. Vegetation type is probably not important, just the presence of it, while soil structure simply has to be such that it holds water to some degree and (perhaps) provides enough small sand grains for mosquito larval hygiene needs. Perhaps there is no dependency of floodwater *Aedes* (or RFVF incidence) on vegetation type or soil structure, just an arbitrary geographical association with the presence of ground-water.

This syntaxonomic study plus the previous one [20] attempt to identify, name syntax, and categorize the changes within and between wetlands and depressions. Various RVF studies detail environmental conditions including meteorology, climatology [15], mention wetlands, depressions [21] and vegetation [17] Rossouw *et al*., 2021, but do not differentiate between wetland categories, or describe and define vegetation communities compiled in formal phytosociological studies.

Vegetation structural diversity can be remotely measured via satellite to ascertain NDVI [15] as well as using LiDAR (Lang *et al*., 2023), with both methods being able to ascertain α diversity. To date, this and previous papers have concentrated on vascular plant diversity but not considered the soil microbiome diversity. As shown by Lang *et al*., (2023), it may the most important factor to explain periodic outbreaks that occur over decades, time sufficient for the soil microbiome, including mycorrhizae and bacteria, to recover.

### Ecological context for RVFV vectors

The ability of the RVFV to survive in wetlands for extended periods remains a hypothesis [25]. Soil and wetland plant species are likely two common factors providing suitable endorheic pan habitat for *Aedes*, but the exact environmental factors are still under investigation.

Low-level RVFV transmission has been demonstrated in the Free State by the presence of unvaccinated seropositive livestock that were born after the 2010-2011 outbreaks [51], with an isolated RVF outbreak on May 2018 that occurred within our study area [52]. Given the temperate climate in the Free State, RVFV must overwinter within the mosquito population and simulations suggest that transovarial transmission plays an important role in supporting this (Rostal *et al*., 2025). Gargan *et al*. (1988) [21] cited evidence (Alexander 1957) that floodwater mosquito eggs could survive in wetland soil and accompanying vegetation, during lengthy, dry periods, and when the infected eggs were hatched, the resultant adult mosquitoes could transmit RVFV. Under the cryptic transmission hypothesis (Linthicum *et al*., 2016; Njenga *et al*., 2019), circulation of RFV is maintained within or between wild and domestic ruminants and RVFV vectors.

Infected female floodwater *Aedes* mosquitoes lay infected eggs in wetlands, which, under the right conditions can initiate focal outbreaks which may lead to large RVF outbreaks, such as seen in SA in 1950 - 1951, 1974 - 1975 and 2010/11 [1, 50, 53]. However, we did not detect RVFV infection in the mosquitoes in this study (Gregory *et al.*, In prep).

This is the second wetland vegetation study in South Africa that links RFV ecology and vegetation science. The study includes detailed B-B surveys, as well as an extensive line-point vegetation density survey. It is in situ, ‘trough-to-ground’ related to NDVI satellite maps, detailed field work to allow future correlation between vegetation associations, vegetation density, mosquito numbers per site, and, which can be combined with findings from the study-data by Anyamba *et al*. (2020) using, geographically large-scale, satellite-derived climatology, meteorology, rainfall, land surface temperature (LST), and normalized difference vegetation index (NDVI) data to understand both broad and detailed environmental conditions for RFV.

The peaks for rainfall, land surface temperature (LST), vegetation cover and mosquito numbers need to coincide. What is shown with the Anyamba *et al*. (2022) [15] study, is a shift in to later in the year of rainfall, LST using NDVI as a surrogate and wetland plant growth, which may not coincide to produce the most suitable breeding conditions for *Aedes*. What has not been included in these models are soil conditions, particularly the microbiome diversity and how quickly it recovers which may affect vegetation recovery and growth, after extended drought periods, to which the region is frequently subject.

### Future Research

Further research may include surveying for floodwater *Aedes mcintoshi* egg cases to investigate whether there is a specific association with one or more specific wetland plants or show the link between wetland vegetation communities and the potential for RVFV transmission. Some of this complex labour-intensive work has been done in Australia (Dale & Knight, 2008), but it has not yet been done in sub-Saharan Africa. Laboratory experiments that attempt to rear floodwater *Aedes* mosquitoes and demonstrate or disprove transovarial transmission under controlled conditions area needed, but have been difficult to initiate because of problems with laboratory colonization of the vector and a shortage of suitableACL-3 (arthropod containment level 3) and higher containment facilities in Africa.

Management of RVFV and control of outbreaks are currently done via large-scale vaccination campaigns that are costly and difficult to organise, or by insecticide spraying or larvicide application of potential mosquito breeding sites [54]. Pesticide use is less effective and are *ad hoc* methods with detrimental impacts on wetland water quality, animal and plant species. A more detailed understanding of the wetland vegetation and broad wetland ecology would allow for more specific and cheaper management practice, as well as having a more precise targeted approach.

### Climate Change

Wetlands are vitally important azonal ecosystems that are also highly threatened by agriculture through grazing for animals, crop farming and drinking water. All these anthropogenic forces are transforming and eliminating wetlands [32]. An additional factor that will impact the future ecology of wetlands is climate change [55]. It is unknown what the influence of climate change will be on the *Aedes* habitat as suitable breeding sites and therefor the risk of RVF outbreaks. With increased storm intensity and greater rainfall, climate change [55], may possibly increase the occurrence of RVF outbreaks. This makes it all the more important to identify wetland vegetation habitat and the overlap with breeding distributions of *Aedes* floodwater mosquitoes (Dale & Knight 2008). As ecosystem services wetlands provide changes, it is important to establish baseline estimates of biodiversity, mosquito abundance, water quality and vegetation cover and abundance. All of these ecological parameters will be influenced by increased temperatures. Inland wetlands in South Africa are the habitat for *Aedes* mosquitoes, which are believed to be responsible for RVF outbreaks that cause human and animal deaths and severe economic loss [56].

Beyond wetland characterization we provide in here, there remain important questions regarding the association between the ecological factors presented here and the maintenance of RVFV transovarially transmitting vectors. We suggest that future research focus on:

1. What are the critical climatological and local meteorological conditions that drive the presence and abundance of floodwater *Aedes* and potentially future outbreaks?
2. Is there sufficient circulation of RFV virus in domestic and possibly wild animals [51], to maintain RVFV during non-outbreak periods of 20-plus years?
3. Are there important associations between suitable wetland types for breeding RVFV vectors and specific vegetation patterns {Linthicum, 1999 #40;Brand, 2018 #8}? Or is it a complex mix of vegetation, soil type, structural soil and soil microbial diversity (Lang *et al*., 2023).

## Conclusion

Our work provides critical insights into wetland vegetation composition in areas experiencing historical RVFV outbreaks.

After severe droughts as in 2015/2016 we report wetland vegetation changes potentially driven by soil removal subsequent overgrazing, which could deplete the soil mycorrhizae and bacteria diversity. Despite this, we continue to find floodwater *Aedes* mosquitoes at these sites, suggesting they remain viable breeding sites. We do not know whether RVFV continues to survive within these populations of floodwater *Aedes*.

Whether extensive livestock and human exposures to RVF virus are associated with transovarially infected eggs or the sporadic introduction of RVF virus from different endemic area or by endemic, cryptic transmission cycles, the ecology of wetlands in the temperate inland grasslands is critical to amplification of RVFV during outbreaks. These habitats provide suitable conditions for maintaining large populations of suitable vectors of RVFV. The same Grassland and temperate Savanna biomes provide for productive livestock farming, particularly of sheep which are highly susceptible to infection with RVF virus. Intensive livestock farming serves as a means for virus transmission to be amplified to epidemic proportions. This study has characterized the ecological context for these wetlands by detailed description of the specialised vegetation, the anaerobic soils and the relative abundance of floodwater *Aedes* mosquitos. Canonical understanding is that vertical transmission is the key to maintaining transmission in interepidemic periods in the infection cycle. If proven true, the key that may linking infections in human and other mammals animals would be the infected eggs of floodwater mosquitoes that require specific conditions to hatch and produce swarms of infected mosquitoes {Lumley, 2017 #41}.

## Acknowledgements

Leslie Powrie (SANBI Fellow, National Biodiversity Assessment, Kirstenbosch Research Centre, South African National Biodiversity Institute, Private Bag X7, Claremont 7735, South Africa), for his GIS expertise, time, effort and patience with several virtual meetings, with all the intricacies and layers to produce the excellent vegetation map (Figure 1), showing the sites, wetlands, rivers and major towns in the study area.

Grateful thanks for the time, effort and collaboration given by Linde de Jager for identification of 66 plant species at the University of the Free State Geo Pots Herbarium (BLFU, University of the Free State), Complete and confirm the identification of 144 plant species and entering these data into both the field data books and into the electronic Google spreadsheet. Providing full descriptions of a further 48 specimens from the eight new study sites. And for completing all data records to ExecuVet for scanning and safe keeping, and to ensure all samples are submitted to the UFS herbarium.

All photographs courtesy Robert Brand, Etienne Leroux, Herman Zwiegers, EcoHealth Alliance and ExecuVet.

## Authors Contributions

conceptualisation; methodology; data collection; sample analysis; data analysis; validation; data curation; writing – the initial draft; writing – revisions; student supervision; project leadership; project management; and funding acquisition.

Brand, RF. Writing – original draft, Conceptualisation, Visualization, Methodology, Investigation, Formal analysis, data curation, Field staff supervision, project leadership, project management.

Rostal, M. Writing – editorial draft, Investigation, funding acquisition, project leadership, project management, Formal analysis.

de Jager, L., Plant identification, Sample analysis, data analysis, validation, data curation, Manuscript editing.

van Staden, L., Vegetation Data base entry, data curation, manuscript editing.

Kemp, A., Vector Data collection, sample analysis, data analysis, validation, data curation, project management.

Van Huyssteen, C.W. was responsible for the siting, description, analyses, and interpretation of the soil profiles and data; as well as editing of the manuscript.

Gqalaqha, N., soil collection, data analysis, manuscript review.

Gregory, N. Manuscript revisions, Endnotes, modifications of Figures 4, 5, 6 and 7. Anyamba, A., provided climate data, interpretation, and revisions

Cordel, C., Project management, Project Leadership, Data Curation, staff management, manuscript revision.

A., Karesh, W.B., Project Leadership, Project Management, Funding acquisition, Manuscript revision.

Paweska, J.T. Project Management, Project Leadership, Student Supervision, funding acquisition, revision and editing of the manuscript.

## Supplementary Material

Supplement 1/Appendix A. Full Syntaxonomical Table for Wetland Vegetation for Rift Valley Fever outbreak.

Supplement 2/Appendix B. Species list of plants with authors names and botanical voucher numbers.

Supplement 3/Appendix C. Full vegetation description of all 12 communities, 18 sub-communities and 10 variants.

Supplement 4/Appendix D. Theoretical and Practical Considerations for Line-point Field work.

